# *Bordetella* Spp. Utilize T3SS To Promote Secretion of IL1RA Leading to Long-Term Persistent Infections

**DOI:** 10.1101/2022.09.26.509564

**Authors:** Tyler L. Williams, Connor Roan, Nicholas First, Sarah Johnson, Monica C. Gestal

## Abstract

A common feature of pathogens is their ability to suppress host immune responses. Understanding the molecular mechanisms and the common pathways that bacteria utilize to block host immune signaling cascade might provide novel avenues for vaccine and therapeutic development. Preventable infectious diseases remain one of the major causes of morbidity and mortality worldwide and the current rise in antibiotic resistance is increasing this burden. *Bordetella* spp. are respiratory pathogens that cause the long-term illness known as whooping cough. *Bordetella* infections cause over 150,000 deaths each year, despite a vaccine being available. In our studies, we used the mouse pathogen *B. bronchiseptica* to investigate the pneumonic stage of disease, which very well mimics the fatal disease caused by *B. pertussis*. In our previous work we discovered a *B. bronchiseptica* mutant, RB50Δ*btrS*, that clears rapidly from the lungs of mice and generates protective immunity that lasts for at least 15 months post-challenge. Combining the mouse immunological tools and both bacteria, the wildtype RB50 and mutant RB50Δ*btrS*, which persists for up to 56 days and clears in 14-21 days, respectively, we investigated the mechanisms by which the wildtype *B. bronchiseptica* blocks host immune response to cause long term lung infection. Previous research indicated eosinophils as critical for rapid clearance of the mutant bacteria from the lungs. *In vitro* assays with eosinophils demonstrated that the RB50Δ*btrS* mutant strain promotes the secretion of pro-inflammatory signals. In contrast, infection of eosinophils with RB50 promoted the secretion of anti-inflammatory signals such as IL1RA. Interestingly, IL1RA was also increased in the lungs of mice infected with the wildtype but not with the mutant RB50 strain. Infection with RB50Δ*bscN*, which lacks a functional type 3 secretion system (T3SS), was sufficient to prevent IL1RA induction suggesting that the bacterial effector responsible for IL1RA upregulation is a substrate of the T3SS. Supplementation with IL1RA after infection with RB50 or RB50Δ*btrS* resulted in increased lung bacterial burden for both bacterial strains. However, more rapid clearance of RB50 was observed after infection of mice in which IL1RA was knocked out. Furthermore, anti-IL1RA antibody treatment promoted rapid clearance of not only RB50 but also the human pathogens *B. pertussis* and *B. parapertussis*. This suggests that IL1RA may be a promising therapeutic target to treat severe cases of whooping cough.

Overall, this work demonstrates that *Bordetella* spp. induces IL1RA expression to promote persistence using the T3SS. Since other bacteria have also been shown to target IL1RA, this may be a conserved bacterial mechanism to promote host-immune suppression.

## INTRODUCTION

Infectious diseases are a major cause of morbidity and mortality worldwide^1^. Due to the increase in antibiotic resistance and parental unwillingness to vaccinate children, the numbers of previously preventable deaths continue to rise^1^. During evolution, pathogens evolved mechanisms to block host immune responses to prevent rapid clearance and allow for reinfection. Understanding the molecular mechanisms that bacteria utilize to suppress host immune responses and prevent the generation of protective immunity can provide novel avenues for development of improved vaccines and therapeutics. In this work, we used the *Bordetella bronchiseptica* murine model to determine how these highly successful pathogens suppress host-immune responses. Here, we identify IL1RA as the mechanism by which *Bordetella* spp. promote long-term persistence and this could be exploited in medical treatments against Bordetella spp. and possible other infections caused by bacterial pathogens that have a functional type 3 secretion system.

*Bordetella* spp. are respiratory pathogens also known for being the etiological agents of whooping cough. Classical *Bordetella* spp. comprises 3 different species; *B. pertussis, B. parapertussis*, and *B. bronchiseptica*, all differing in their host range^2^. *B. bronchiseptica* is the evolutionary ancestor of the other two species, sharing over 98% of the genetic content^3,4^. Moreover, it is a natural pathogen of mice^2,5^ facilitating the study of host immunosuppression at the molecular level. *Bordetella* spp. infection has three main stages; the catarrhal stage, which is characterized by increased mucus secretion; the paroxysmal stage, during which constant violent episodes of cough cause apnea and even loss of consciousness; and the pneumonic phase, which is the main cause of fatal whooping cough^6^. Our work focuses on understanding the molecular mechanisms that *Bordetella* spp. utilize to suppress the generation of robust immune responses during the pneumonic fatal stage of disease. In our previous work, we focused on investigating how *Bordetella* spp. respond to inflammatory signals contained in blood and serum^7^. In response to blood and serum, we observed that *Bordetella* spp. upregulated expression of virulence factors as well as pathways that are known to help bacteria adapt to host microenvironments^7^. Importantly, amongst the different virulence factors, we also identified the increased expression of a sigma factor known as *btrS. btrS* is one of the many regulators of the T3SS^8^, however, it also regulates many other genes involved in virulence, metabolism, and other bacterial phenotypes^9,10^. When studying the effects of *btrS* during the host-pathogen interaction, our results revealed that *btrS* regulates an immunosuppressive pathway that involves multiple known and unknown virulence factors. In the absence of *btrS*, there is a significant increase in the recruitment of B and T cells in the lungs at day 14 post-infection^10^. Mice infected with the *btrS* mutant develop long-lasting protective immunity that prevents reinfection with any of the three classical *Bordetella* spp. including the human pathogens *B. pertussis* and *B. parapertussis*, for at least 15 months post-vaccination^11^. Excitingly, our results revealed that protective immunity was correlated with an increase in the numbers of eosinophils in the lungs^12^.

Eosinophils are known for their function during parasitic infections as well as for their implications during allergic and asthmatic reactions^13^. The fact that eosinophils can kill bacteria has long been known^14^ however it has not gained significant attention until recently, when the role of eosinophil traps during infections, allergic reactions, and asthmatic processes was studied indepth ^15,16^. Eosinophils are critical for immune homeostasis to keep the balance between Th1 and Th2 responses; in fact, they are pivotal to prevent mucosal autoinflammatory disorders^13^. Their anti-inflammatory function is performed through the secretion three main molecules; PDL1 (program death ligand-1), IDO (Indoleamine 2,3-dioxygenase), and IL1RA (IL1 receptor antagonist)^13^. Eosinophils secrete one or more of these molecules to dampen proinflammatory responses and restore the immune homeostatic balance. During *Helicobacter pylori* infections, eosinophils’ expression and secretion of PDL1 dampens pro-inflammatory responses allowing for persistent infections in the gastrointestinal tract^17^. However, this ability of eosinophils to modulate immune responses is not limited to the gut, as eosinophils are also found in other mucosal surfaces where they perform a critical role in maintaining homeostasis. Recent work from our lab demonstrated that during murine infection with a *btrS Bordetella bronchiseptica* mutant, RB50Δ*btrS*, eosinophils are recruited to the lungs. In fact, they are critical for rapid clearance of the infection from the lungs and the generation of adaptive immune responses^12^. Contrary, during infection with the wildtype RB50, eosinophils are not recruited to the lungs and infection persists for at least 56 days, suggesting an immunosuppressive mechanism that might involve eosinophils.

Based on the literature on *H. pylori*^*17*^ and our previous results^12^, we wanted to investigate if the wildtype *Bordetella* spp. promote anti-inflammatory responses that enhance colonization and persistence. Our results indicate that wildtype *Bordetella* spp. promote secretion of IL1RA by eosinophils and epithelial cells by promoting long term persistence. Supplementation with exogenous IL1RA during infection results in increased bacterial burden. Contrary to this, ablation of IL1RA leads to decreased lung colonization. Importantly, our results also revealed that anti-IL1RA can be used as treatment not only against *B. bronchiseptica* but also the human pathogen *B. pertussis*. Overall, our results provide a novel insight into the role of eosinophils during infection as well as a novel immunotherapy that can be used in multiple mucosal infections. We also demonstrated that bacteria promote secretion of IL1RA via T3SS, suggesting that the use of anti-IL1RA therapies can be extended to numerous pathogens that use similar virulence factors.

## RESULTS

### RB50 suppress eosinophil mediated killing

We have previously shown that *Bordetella* spp. can suppress host immune responses via *btrS* mediated mechanism^10^ in an eosinophil dependent manner^12^, suggesting that eosinophils may mediate adaptive mucosal immune responses. It has been shown that eosinophils can phagocytose and kill bacteria^18-20^, and as a result they secrete cytokines that can promote pro- and anti-inflammatory responses^13,21,22^. To investigate eosinophils ability to phagocytose *Bordetella* spp., we differentiated bone marrow progenitors into mature eosinophils following previously published procedures^23^. After confirming >98% differentiation with flow cytometry and/or cytospin, cells were challenged with RB50 or RB50Δ*btrS* at a Multiplicity of Infection (MOI) of 10 for 4 hours to assess bacterial phagocytosis.

Using transmission electron microscopy, we observed that eosinophils can efficiently phagocytized both, RB50 and RB50Δ*btrS* (**Figure 1A**). Interestingly, we observed that bacteria were being phagocytosed by the eosinophils using pseudopodia, in what was not a passive process but rather an active one. The electron microscope photos suggest that the number of bacteria internalized following infection with RB50 or RB50Δ*btrS*, appear to be different. We counted the number of bacteria internalized by eosinophil (**Figure 1B**). Our results suggest that eosinophils challenged with RB50 contained in general between zero and three bacteria, although sometimes they could exceed more than 5 bacteria in the cytoplasm. Interestingly, eosinophils challenged with RB50Δ*btrS* generally contained zero or one bacteria within the cytoplasm, suggesting that either RB50 intracellular survival in eosinophils was greater or that RB50Δ*btrS* was more successfully phagocytosed and killed.

**Figure 1.**
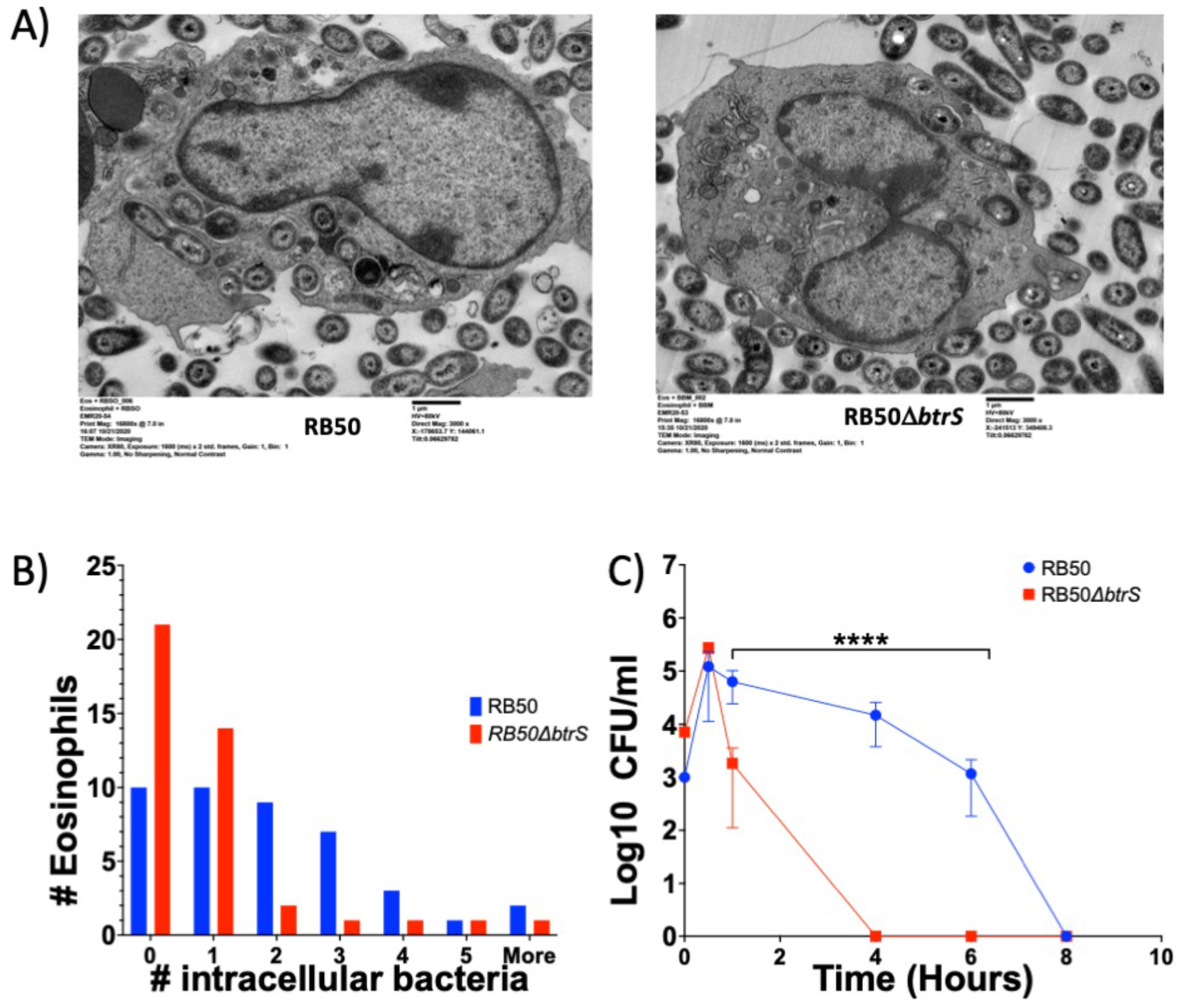
Eosinophils phagocytose and kill *Bordetella bronchiseptica*. (A) Eosinophils were challenged with RB50 (left) or RB50Δ*btrS* (right) at an MOI of 10. At 4 hours post-infection samples were fixed and imaged in the transmission electron microscope. The image is representative of 4 independent experiments done in three technical replicates (n=4×3). (B) Intracellular bacteria were enumerated from at least 50-100 individual eosinophils. We did exclude death eosinophils. Bars represent in blue RB50 and red RB50Δ*btrS*. The x-axis shows number of intracellular bacteria. (C) Eosinophils were challenged with RB50 (blue) or RB50Δ*btrS* (red) at an MOI of 1. This experiment was performed in three individual biological replicates each of them containing 6 technical replicates. At different times post-infection samples were plated to enumerate bacteria and evaluate eosinophil killing. One-way ANOVA was performed **** p<0.0001

To answer this question, we performed a killing assay. We inoculated eosinophils (98% differentiation rate) with an MOI of 1 of either RB50 or RB50Δ*btrS* and at different times post-challenge and bacterial colonies were enumerated (**Figure 1C**). Our results revealed that while RB50 is completely phagocytosed and killed by eosinophils in a period of at least 8 hours, RB50Δ*btrS* is killed in half of the time (4 hours), suggesting that the lower number of Rb50Δ*btrS* could indicate that it is more efficiently phagocytosed and killed by eosinophils. Overall, our results indicate that both bacteria, RB50 and RB50Δ*btrS* can be phagocytosed by eosinophils. Moreover, the results also indicate that RB50Δ*btrS* is killed more efficiently than the wildtype RB50.

### RB50 suppress eosinophil mediated pro-inflammatory responses

Eosinophils contain many cytokines and chemokines in their granules that can orchestrate host immune responses^24^. Due to the differences observed on the interactions between eosinophils and wildtype RB50 or RB50Δ*btrS*, we hypothesized that the cytokine signaling triggered by the RB50 or the RB50Δ*btrS* mutant could be different. We infected eosinophils (98% differentiation rate) with either RB50 or RB50Δ*btrS* at an MOI of 10 and we investigated the cytokine secretome at 4 hours post-infection using the supernatant of our infected eosinophils. The results revealed that infection of eosinophils with RB50 promote an increase in the secretion of IL9 (**Figure 2A**) and IL4 (**Figure 2B**) at two hours post-infection compared with the uninfected control. However, eosinophils challenged with RB50Δ*btrS* did not. Conversely, looking at pro-inflammatory cytokines, eosinophils challenged with RB50 did not increase the levels of secretion of IL2 and IL17 (**Figure 2C, 2D, and 3E**). However, infection with RB50Δ*btrS* significantly increased secretion of IL2 (**Figure 2C**) at only at 2 hours post-infection. At 4 hours post-infection, RB50Δ*btrS* infected eosinophils presented an increased secretion of not only IL2 (**Figure 2D**) but also IL17 (**Figure 2E**). Overall, eosinophils challenged with RB50 wildtype *B. bronchiseptica*, secrete IL9 and IL4 which are cytokines generally associated with Th2 immune responses. Conversely, eosinophils challenged with RB50Δ*btrS* secrete pro-inflammatory cytokines that promote Th1/Th17 responses.

**Figure 2.**
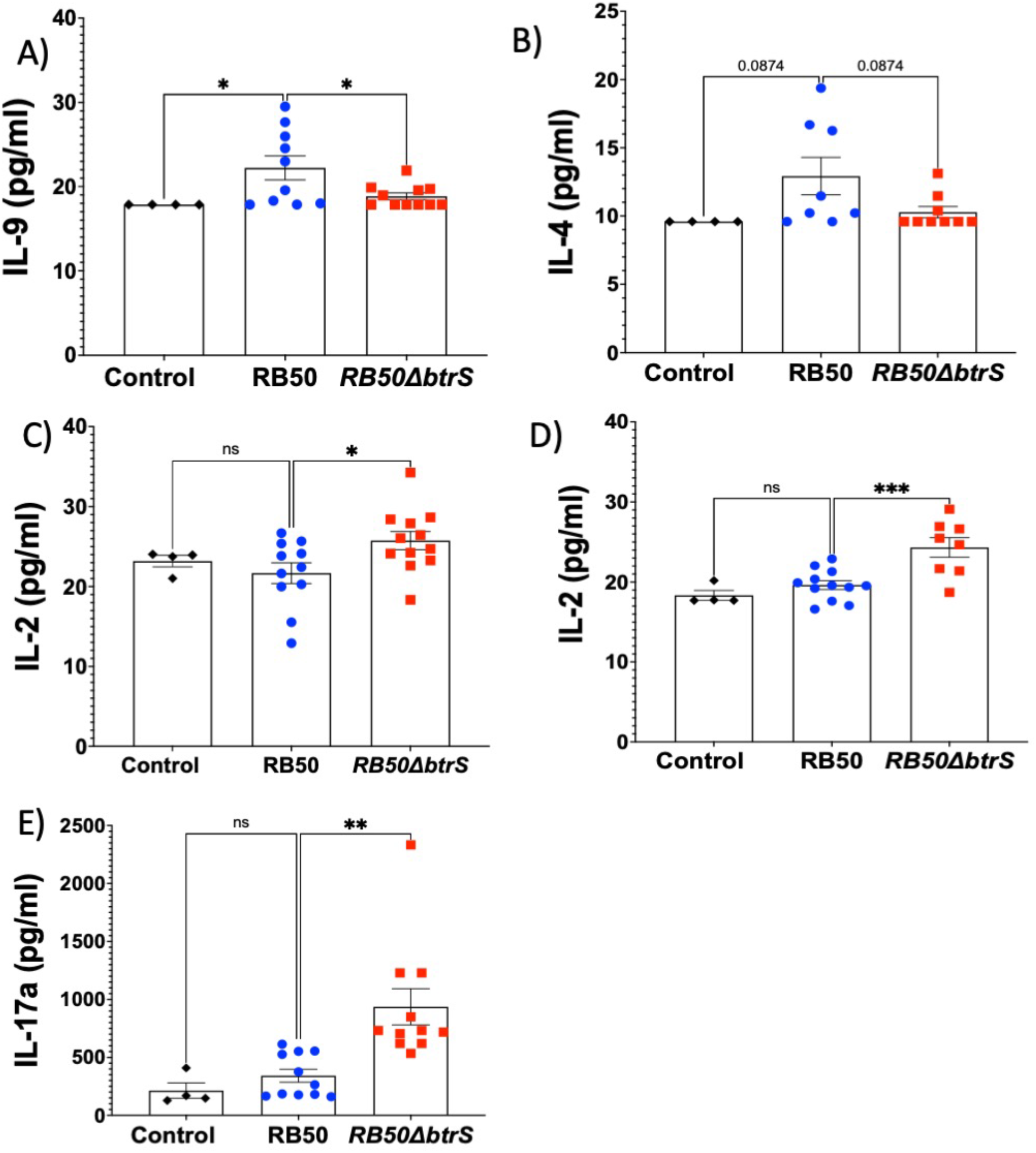
*Bordetella bronchiseptica* blocks eosinophil pro-inflammatory responses via *btrS* mediated mechanism. Eosinophils were unchallenged (rhomboid black) or challenged with RB50 (circle blue) or RB50Δ*btrS* (square red) at an MOI of 10. At 2- and 4-hours postinfection the supernatant was collected to performed a LegendPlex analysis. This experiment was performed in 4 individual biological replicates each one containing 3 technical replicates. Each symbol corresponds with the average of two technical replicates in the ELISA assay. (A) Bars shows pg/ml of IL9 measured at 2 hours post-infection. (B) Bars shows pg/ml of IL4 measured at 2 hours post-infection. (C) Bars shows pg/ml of IL2 measured at 2 hours post-infection. (D) Bars shows pg/ml of IL2 measured at 4 hours post-infection. (E) Bars shows pg/ml of IL17a measured at 2 hours post-infection. One-WAY ANOVA was performed to analyzed the data. * p<0.05, ** p<0.01, *** p<0.001, ns = non-significant. The values on (B) indicate the p-value.

### RB50 promotes IL1RA expression in eosinophils

It is known that eosinophils prevent auto-inflammatory disorders on the mucosal surfaces, such as inflammatory bowel syndrome in the guts^25,26^ by secreting Interleukin-1 receptor antagonist (IL1RA)^27-29^, Program death ligand 1 (PD-L1)^17,22^, and Indoleamine 2,3-dioxygenase (IDO)^13,30^. Based on the differences identified in the cytokine profile, we hypothesized that RB50 might promote the expression of one of these molecules, to suppress the secretion of IL17 by eosinophils. Based on previous literature we believe that we will identify an increase in IL1RA that will be responsible for suppressing IL17 secretion in eosinophils challenged with RB50^28^.

Using a minimalistic approach co-culturing bone marrow derived eosinophils (98% differentiation rate) challenged with both bacteria at an MOI of 10, we found that infection with RB50 but not RB50Δ*btrS* promotes increase expression of mRNA levels of IL1RA by eosinophils (**Figure 3A**). No changes in the expression of IDO (**Figure 3B**) and PD-L1 (**Figure 3C**) were detected after infection with RB50 or RB50Δ*btrS* compared with the uninfected control. Thus, these results suggest that RB50 promotes IL1RA expression in eosinophils, but not PD-L1 or IDO, indicating that RB50 promotes expression of IL1RA via a *btrS* mediated mechanism.

**Figure 3.**
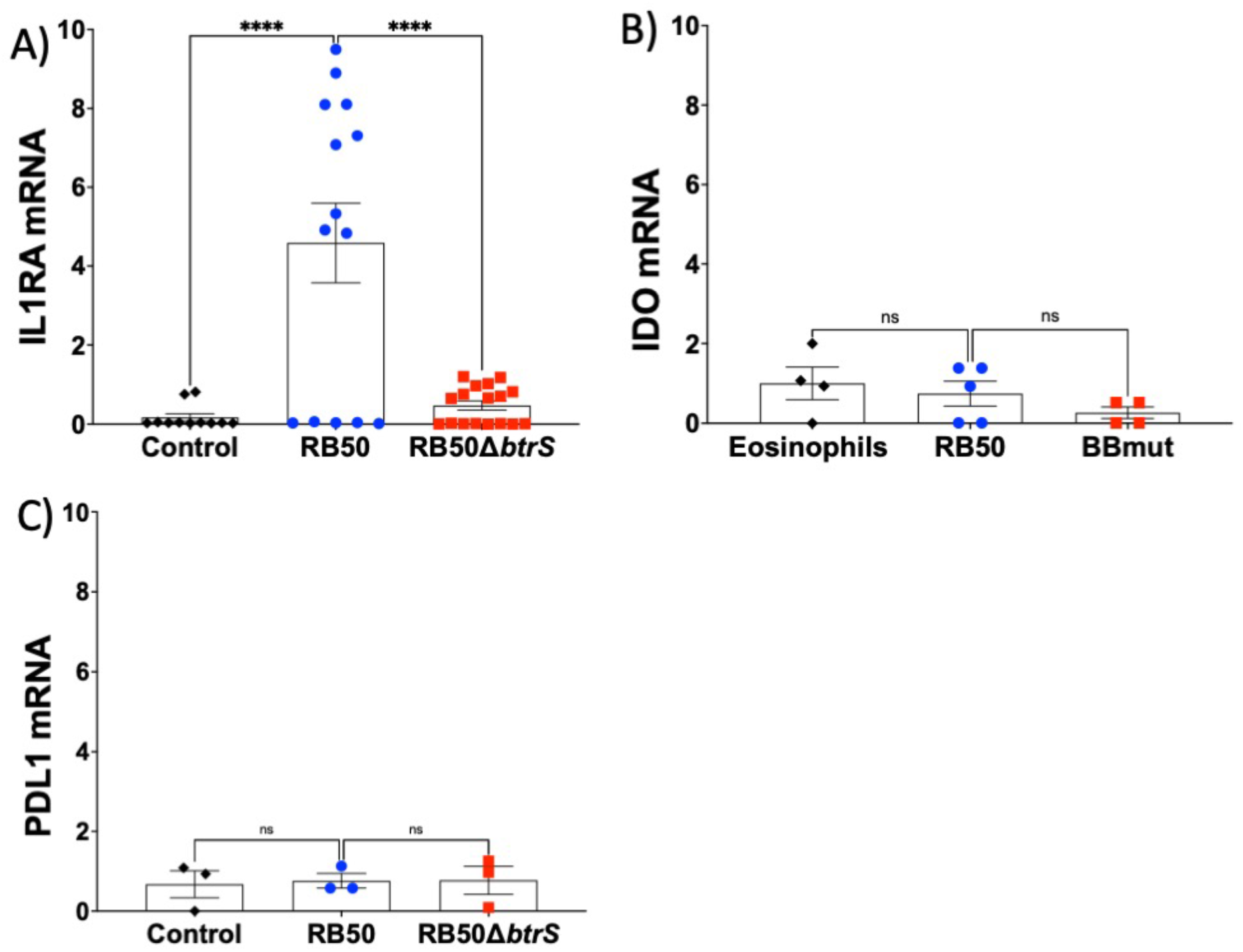
*Bordetella bronchiseptica* promotes secretion of IL1RA by eosinophils via *btrS* mediated mechanism. Eosinophils were unchallenged (rhomboid black) or challenged with RB50 (circle blue) or RB50Δ*btrS* (square red) at an MOI of 10. At 4 hours post-infection RNA was extracted to perform qRT-PCR. ΔΔct was normalized to actine. This experiment was performed in 3 individual biological replicates each one containing 3 technical replicates. Each symbol corresponds with the average of the three technical replicates in each qRT-PCR assay. (A) Bars shows mRNA levels (fold change) of IL1RA at 4 hours post-infection. (B) Bars shows mRNA levels (fold change) of IDO at 4 hours post-infection. (C) Bars shows mRNA levels (fold change) of PDL1 at 4 hours post-infection. One-WAY ANOVA was performed to analyzed the data. **** p<0.0001, ns = non-significant.

### RB50 promotes expression of IL1RA in vivo

It has been previously shown that IL1RA has a role in vivo by aggravating infections^31^, one example being *Staphylococcus aureus* septicemia, where it has been shown that pathology is increased when mice are treated with IL1RA^32^. In addition, IL1RA has been shown to increase during infections with some bacteria^31,33^. One example of IL1RA increasing is in cystic fibrosis patients infected with *Pseudomonas aeruginosa*^34^.

To evaluate the effects of RB50 and RB50Δ*btrS* infection on IL1RA, PDL-1 and IDO *in vivo*, we challenged mice with PBS, RB50, or RB50Δ*btrS*. At different times post-infection, we extracted RNA from the lungs to perform qRT-PCR and evaluate the changes in mRNA levels for IL1RA, IDO and PDL1 using actin to normalize the results. Following infection with RB50, the levels of IL1RA mRNA increase as early as day 1 post-infection, peaking at day 7 and then returning to basal levels at day 14 after infection (**Figure 4A**), showing a similar trend to that previously reported for *P. aeruginosa* infection^34^. Interestingly the mRNA levels of IL1RA barely changed after infection with RB50Δ*btrS*. Similar to our results *in vitro*, no changes on mRNA levels for IDO (**Figure 4B**) nor PDL1 (**Figure 4C**) were detected after infection with RB50.

**Figure 4.**
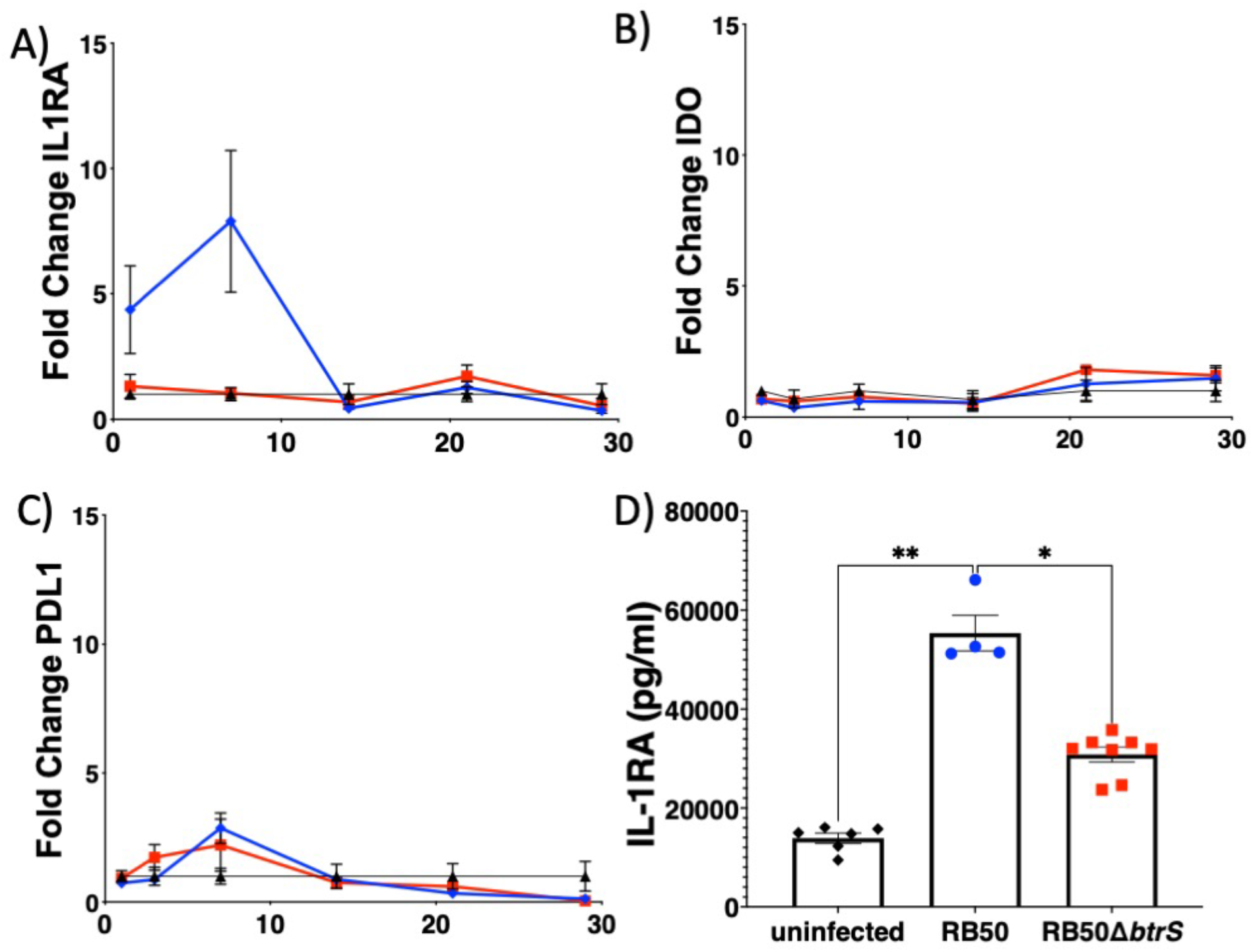
*Bordetella bronchiseptica* promotes secretion of IL1RA in vivo via *btrS* mediated mechanism. Balb/c mice were unchallenged (rhomboid black) or challenged with 30μl of PBS containing 5×10^5^ RB50 (circle blue) or RB50Δ*btrS* (square red). At different post-infection times lungs were collected to extract RNA and qRT-PCR was performed. ΔΔct was normalized to actine. This experiment was performed in 2 individual experiments each one containing 3 mice of each group. Each symbol corresponds with the average of the three technical replicates in each qRT-PCR assay. (A) Lines and symbols show the average and SEM of mRNA levels (fold change) of IL1RA. (B) Lines and symbols show the average and SEM of mRNA levels (fold change) of IDO. (C) Lines and symbols show the average and SEM of mRNA levels (fold change) of PDL1. (D) At day 7 post-infection, lungs were collected in PBS and protease inhibitor. After homogenizing the supernatant was used to evaluate IL1RA ELISA. Each symbol represents the average of two technical replicates and the bars show the pg/ml of IL1RA secreted in lungs. One-WAY ANOVA was performed to analyzed the data. * p<0.05; ** p<0.01.

To confirm that the changes in mRNA levels were related to differences in the protein levels, an ELISA using (with lung homogenized with protease inhibitors) was performed using lungs of infected mice at day 7 post-infection. This was the time that coincides not only with the peak of infection but also the peak in mRNA levels of IL1RA observed. Our results show that concentration of IL1RA in the lung homogenate increased at day 7 post-infection in mice challenged with RB50 (**Figure 4D**). Mice challenged with RB50Δ*btrS* also presented a small increase but that was not comparable to that shown with the wildtype strain. Thus, our results revealed that infection with the wildtype bacteria RB50 promotes increased expression and secretion of IL1RA, an immunosuppressive molecule, that has the potential to enhance colonization and facilitate long term persistence.

### IL1RA promotes persistence in vivo

Previous literature associate supplementation with IL1RA in patients that receive treatment with anakinra and a worse outcome during infectious diseases, mostly correlated with increased bacterial burden^32,35^. Due to the difference in persistence in lungs observed following infection with RB50 or RB50Δ*btrS*^10^, we hypothesized that IL1RA might facilitate persistence. To determine the effects of IL1RA on length of infection with RB50 and RB50Δ*btrS*, we used a knockout mouse model (Ilr1n^-/-^)^31,36^ combined with the wildtype C57BL/6J. We evaluated lung colonization levels at day 14 post-infection with the RB50 and RB50Δ*btrS*. We found that in the absence of IL1RA in the knockout mice bacterial burden of RB50 in the lungs was a decreased by 3 logs (Fig 5A). No differences were observed on mice infected with the RB50Δ*btrS* mutant comparing Il1rn^-/-^ and C57BL/6J mice. This was expected as by day 14, infection with RB50Δ*btrS* is nearly cleared from the lungs.

**Figure 5.**
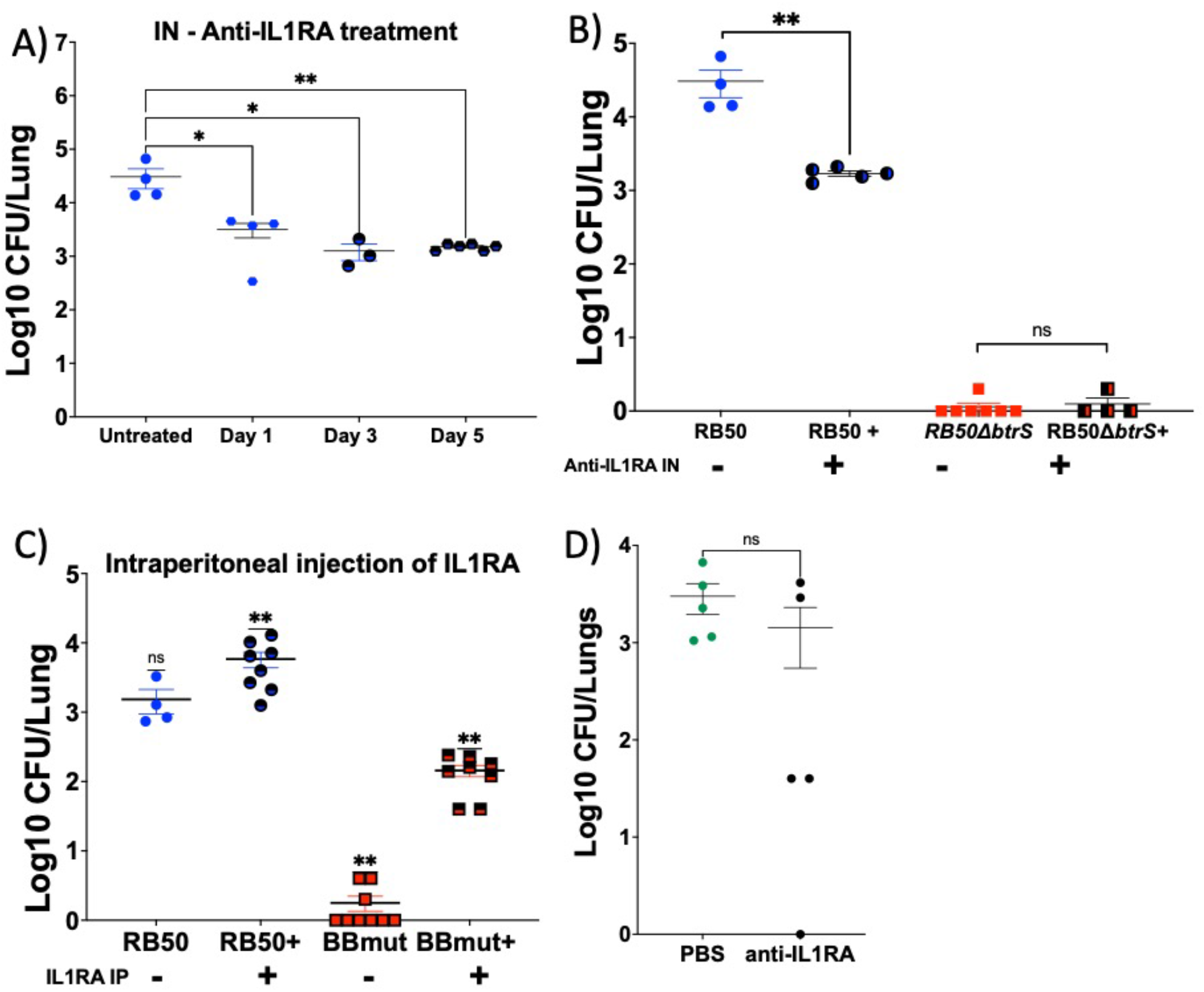
IL1RA promotes *Bordetella bronchiseptica* persistence in the lungs. Balb/c mice were unchallenged (rhomboid black) or challenged with 30μl of PBS containing 5×10^5^ RB50 (circle blue) or RB50Δ*btrS* (square red) in two independent experiments containing between 2-4 mice each time. (A) At day 1, 3 or 5 post-infections, mice started treatment with anti-IL1RA antibodies until day 14 post-infection. Lung bacterial burden was enumerated. Symbols log10CFU of RB50. (B) At day 5 post-infection mice started daily treatment with intranasal anti-IL1RA. At day 14 bacterial burden was enumerated in the lungs. (C) At day 1 post-infection mice started daily treatment with intraperitoneal IL1RA. At day 14 post-infection bacterial burden was enumerated in the lungs. (D) Balb/c mice were intranasally infected with *B. parapertussis*. At day 5 we initiated daily treatments with anti-IL1RA up to day 14 post-infection (black). The other group was infected and untreated (green). At day 14 lung bacterial burden was enumerated. One-WAY ANOVA was performed to analyzed the data, except figure (C and D) were T-test Wilcoxon was performed. * p<0.05; ** p<0.01; ns = non-significant.

To further confirm that the effects of IL1RA on bacterial persistence were due to the absence of IL1RA and not another reason, we used anti-IL1RA antibodies intranasally every 24 hours. We first evaluated when to start the treatment, so we infected mice with RB50 and started treatment at day 1, 3, and 5 post-infections (**Figure 5A**). We intranasally treated them daily until 14 and following euthanasia, we enumerated bacterial colonies in the lungs. We found that treatment with anti-IL1RA significantly decreased bacterial burden in the lungs. Interestingly, no differences were found in regards as to when to start the treatment. Starting treatment at day 1, 3 or 5 post-infections (**Figure 5A**) all rendered the same reduction in the lungs bacterial burden. Based on these results we decided to start treatment at day 5 post-infection for the following experiments, as this day is the day at which whooping cough patients most commonly report to the doctor.

We infected mice with RB50 and RB50Δ*btrS* and provided treatment from day 5 up to day 14 and subsequently evaluated bacterial colonization in the lungs (**Figure 5B**). Our results reveal that treatment with anti-IL1RA antibodies significantly reduced bacterial burden following infection with RB50. However, similar to the results obtained with the IL1RA knockout mice, we did not observe decreased burden after infection with RB50Δ*btrS*, possibly because by day 14 the infection is already cleared.

We then wanted to investigate if supplementation with IL1RA will promote increased bacterial persistence. To answer this question, we challenged mice with RB50 and RB50Δ*btrS*. Twenty-four hours post-infection we began daily treatments with intraperitoneal injections of IL1RA until day 14 post-infection. Our results revealed that supplementation with IL1RA leads to an increase in bacterial burden during infection with RB50 and RB50Δ*btrS* (**Figure 5C**), supporting previous observations that IL1RA supplementation increases bacterial burden^37^. Overall, these results indicate that IL1RA promotes bacterial persistence in the lungs. Contrary, ablation of IL1RA using a knockout mouse model or antibodies anti-IL1RA, results in more rapid clearance of *B. bronchiseptica* infections.

### Anti-IL1RA antibodies promote rapid clearance

Based on our previous results indicating that anti-IL1RA antibodies lead to decreased bacterial burden, we wanted to evaluate if anti-IL1RA antibodies could be used to promote clearance of infection by the three classical *Bordetella* spp. Groups of mice infected with the RB50 wildtype strain of *B. bronchiseptica*. While we did not reach statistical significance to more clearance of BP536 *B. pertussis*^*7,11*^, and BPP12822 *B. parapertussis*^*7,11*^, treated intranasally every day after day 5 post-infection with 10μl of PBS or 10μl of PBS containing 100ng of anti-IL1RA antibodies. Our results clearly reveal a trend of less bacterial burden after treatment, and in our future studies we will be testing different dosages and concentrations (**Figure 5D**). Overall, our results suggest that anti-IL1RA antibodies could be investigated as a treatment for *Bordetella* spp. infections.

### The bacterial type 3 secretion system (T3SS) is required for IL1RA expression

Our data indicates that Bordetella spp. increase levels of IL1RA during infection to facilitate persistence. Moreover, treatment with anti-IL1RA antibodies promotes rapid clearance. To follow these findings, we investigated the bacterial mechanisms by which RB50 promotes IL1RA, while RB50Δ*btrS* does not. *btrS* is a sigma factor that controls the expression of many known and unknown virulence factors including the type 3 secretion system (T3SS)^8-10^. T3SS is one of the most important bacterial virulence factors that many pathogens possess, known for promoting anti-inflammatory responses. In fact, during *Bordetella* spp. infections T3SS promotes an increase in IL10 after infection to facilitate colonization and persistence^38^. Based on these data, we hypothesized that T3SS might promote the expression and secretion of IL1RA as one of the multiple mechanisms by which the T3SS promotes anti-inflammatory responses.

To test that hypothesis, we challenged mice with a T3SS mutant, RB50Δ*bscN*, which lacks the ATPase that allows the T3SS to pump effectors out of the bacteria cell^39^. An important characteristic of this RB50Δ*bscN* mutant is that it clears more rapidly than the RB50^40^, although not as fast as the RB50Δ*btrS*, providing a good candidate to test our hypothesis. We infected mice with RB50, Rb50Δ*btrS*, and RB50Δ*bscN*, at day 7 post-infection we evaluated levels of expression of IL1RA in the lungs of infected mice (**Figure 6A**). Indeed, mice infected with the RB50Δ*bscN* mutant did not present significantly increased levels of IL1RA in the lungs. RB50 revealed an increase in the levels of mRNA of IL1RA, while contrary, RB50Δ*btrS* and RB50Δ*bscN*, did not increased expression of IL1RA compared to untreated controls, indicating that T3SS promotes IL1RA expression.

**Figure 6.**
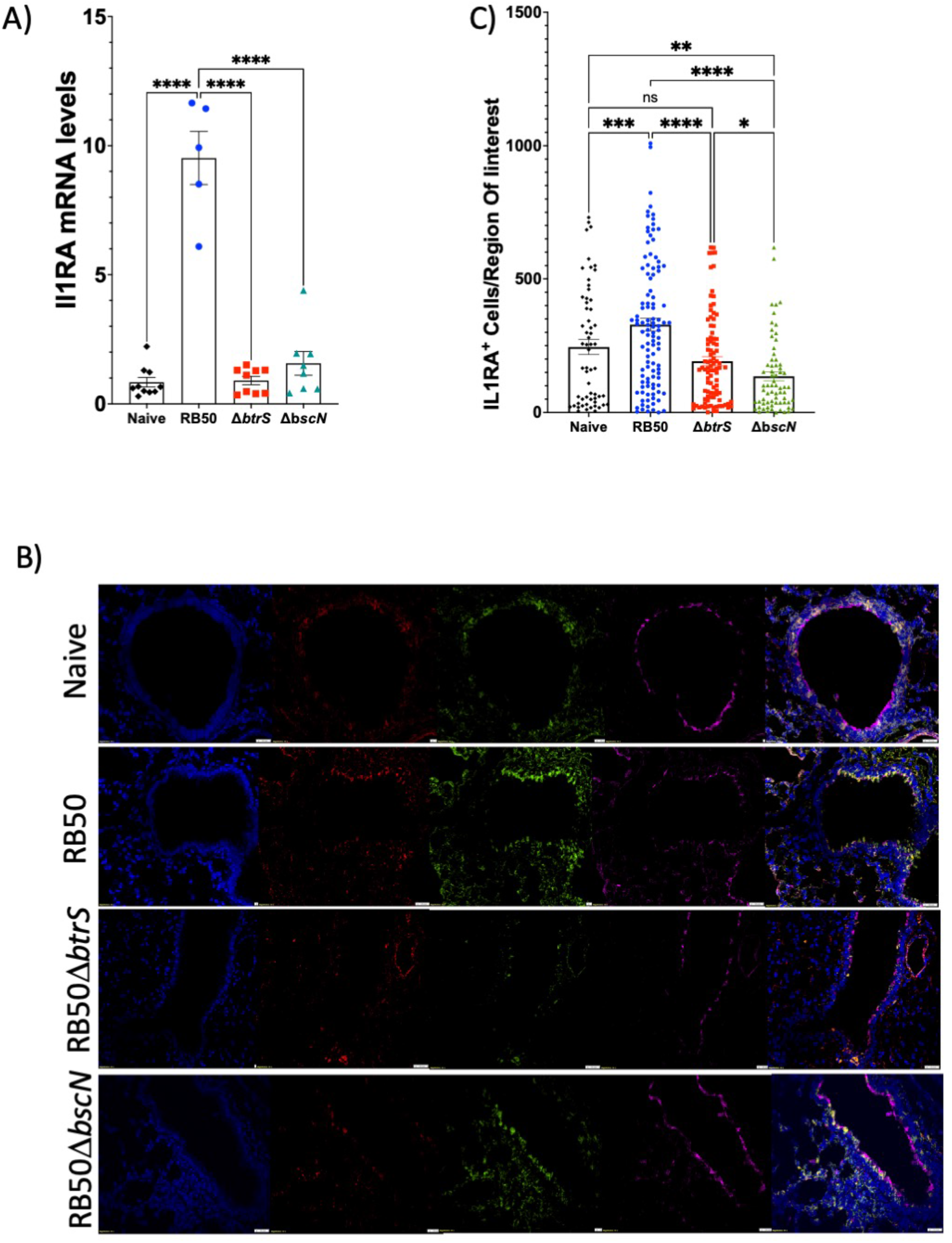
Bacterial T3SS promotes expression of IL1RA during Bordetella bronchiseptica infection. Balb/c mice were unchallenged (rhomboid black) or challenged with 30μl of PBS containing 5×10^5^ RB50 (circle blue), RB50Δ*btrS* (square red), or RB50Δ*bscN* (triangle green) in two independent experiments containing between 2-4 mice each time. (A) At day 7 post-infection, lungs were collected to extract RNA and perform qRT-PCR. ΔΔct was normalized to actine. This experiment was performed in 2 individual experiments each one containing 3 mice of each group. Each symbol corresponds with the average of the three technical replicates in each qRT-PCR assay. Bars represent mRNA levels (fold change) for IL1RA. (B) At day 7 post-infection lungs of 6 different animals per condition were fixed in 4% PFA followed by paraffin embedding, and sectioning. Immunofluorescence staining shows nucleus (blue, Hoechst), epithelial cells (red, Epcam), and IL1RA (green, anti-IL1RA). (C) Six Balb/c mice, three section of each were stained. Using the Keyence microscope, 10 areas of each section were acquired and the value of positive IL1RA staining per region of interest was evaluated using their software. Bars represent positive IL1RA signal per region of interest. One-WAY ANOVA was performed to analyzed the data, except figure (C) were T-test Wilcoxon was performed. * p<0.05; ** p<0.01; *** p<0.001, **** p<0.0001; and ns = non-significant.

To further confirm the results, immunofluorescence microscopy was performed using paraffin embedded sections of the lungs of uninfected mice as well as day 7 post-challenged with RB50, RB50Δ*btrS*, and RB50Δ*bscN*. We stained the nucleus (blue), IL1RA (green), and epithelial cells using Ep-Cam as a marker (red) (**Figure 6B and 6C**). Our naïve group have certain amount of constitutive IL1RA produced by several cells. These were expected as IL1RA is constitutively produced to maintain homeostasis and prevent auto-inflammatory reactions^28^. The mice infected with the RB50 strain have a significant number of epithelial cells that were highly positive to IL1RA. We believe that although eosinophils might contribute as important mediators of the Il1RA increase, epithelial cells are the major source. RB50Δ*btrS* have lower IL1RA positive signal and it was localized only at the border of the epithelia. This was expected based on our mRNA, protein results and animal experiments. Mice challenged with RB50Δ*bscN* also have lower signal, similar to that obtained with our RB50Δ*btrS* group, confirming the mRNA results previously obtained. Altogether suggesting that the *Bordetella* T3SS promotes IL1RA as part of one of the multiple mechanisms by which T3SS dampen host-inflammatory responses.

## DISCUSSION

Pathogens are characterized for their ability to suppress host immune responses^2^. Most of the known mechanisms of bacterial pathogenesis focus on interactions between bacteria and classical phagocytes or innate immune cells, including macrophages^41-43^, neutrophils^44,45^, and dendritic cells^46,47^. However, other immune cells generally classified as type 2, such as eosinophils’^48-50^ or mast cells^51-53^, which have been historically overlooked during bacterial infections, are gaining attention. In this work we follow up on our previous findings to investigate the mechanism by which eosinophils mediate long term persistence of *Bordetella* spp. in the respiratory tract. Our results reveal that *Bordetella* spp. suppress eosinophil effector functions via *btrS* associated mechanism, promoting survival and eosinophil mediated killing. Moreover, eosinophils secrete IL1RA and this increase of IL1RA is also observable in lungs in vivo. Increased IL1RA promotes long term persistent infection in the lungs. Conversely, anti-IL1RA antibodies decrease lung bacterial burden after infection with any of the other classical *Bordetella* spp. Finally, we identify that the bacterial mechanism to promote IL1RA secretion is via T3SS, one of the most well-known bacterial virulence factors^54^, suggesting that this might be a conserved mechanism to promote immune-suppression.

Eosinophils are granulocytes mostly associated with pathological states of disease such as asthma or allergies^16,48^. However, their role during immune homeostasis is known to be important especially in mucosal surfaces such as the gut, where eosinophils prevent autoinflammatory disorders such as Chron’s or inflammatory bowel syndrome^25,55^. Moreover, their function during cancerous processes is also being investigated as eosinophils play critical roles in the tumor microenvironment, where eosinophils are being proposed as a target for the development of future immunotherapies due to their crosstalk with lymphocytes and other immune cells^56^. However, their role during infectious diseases has been overlooked with research focusing primarily on responses to parasitic infections^57,58^. The fact that eosinophils can phagocytose and kill bacteria has been long known^20^, and their ability to form traps during bacterial infections has been proven critical during infections with *Staphylococcus aureus* and other bacteria^59^. In this work, we demonstrate that eosinophils can efficiently phagocytize and kill *Bordetella* spp. Importantly, we show that eosinophils can more effectively kill a mutant bacterium, RB50Δ*btrS* which cannot suppress host immune response, than the RB50 wildtype, suggesting that RB50 blocks eosinophil mediated killing via *btrS* mediated mechanism. Interestingly, when analyzing the differences in the cytokine profiling we identify that while eosinophils challenged with the RB50Δ*btrS* mutant bacterium promote the secretion of pro-inflammatory cytokines, such as IL17 while on the contrary, eosinophils challenged with the wildtype increase the expression of antiinflammatory cytokines such as IL1RA.

Recent literature demonstrates that eosinophils are utilized by bacteria to promote persistence in the gut^17^ and the respiratory tract^12^. It has been shown that eosinophils in the gut dampen pro-inflammatory responses by secreting PDL1 in response to *Helicobacter pylori* infection facilitating persistence^17^. We have previously demonstrated that infection with RB50DbtrS promotes Th1/Th17 responses^10,11^. Here our results indicate that in the respiratory tract, eosinophils, together with epithelial cells, secrete IL1RA in response to infection with *B. bronchiseptica* as an alternative mechanism to dampen pro-inflammatory responses.

IL1RA blocks IL1Receptor-1, competing with IL1Δ and IL1α for its binding and suppressing the subsequent immune signaling^60,61^. IL1RA is used as a treatment for autoinflammatory disorders, and it is commercially available. Anakinra is used for the treatment of autoinflammatory diseases such as rheumatoid arthritis and others^62^. Patients who receive anakinra (IL1RA) during long periods of time, have higher risk of suffering from infectious diseases^37^. But interestingly, while infections and bacterial burden increase, pathology can be decreased with IL1RA^31,34,63^, highlighting the finely tuned balance that must be modulated during pro-and anti-inflammatory responses. Our results demonstrate that during infection with the wildtype bacteria which persist for up to 56 days in the lungs^10^, there is an increase in the levels of IL1RA at early stages of infection. Conversely, infection with our RB50Δ*btrS* strain, which clears by day 14-21^10,12^, does not present the same increase in IL1RA levels suggesting that IL1RA might mediate the long-term persistence of the wildtype bacterial infection. Similarly, this increase in IL1RA has been previously shown following immunization with acellular pertussis^64^, mRNA vaccines^65^, and infections caused by *Pseudomonas aeruginosa*^*34*^, or *Chlamydia*^*66*^. Thus, suggesting that pathogens might promote an increase of IL1RA to enhance colonization and persistence. We investigated the effects of anti-IL1RA antibodies during bacterial infection and our promising results revealed a decrease in the lung bacterial burden not only during infection with *B. bronchiseptica*, but also the human pathogens, *B. pertussis* and *B. parapertussis*.

Investigating the bacterial mechanism that drives the increase of IL1RA, our results indicate that it is mediated by the bacterial Type 3 Secretion System (T3SS). Previous research on the mechanisms of immune suppression of the T3SS focused on classical innate phagocytes^54^, including macrophages^67^, neutrophils^46^, and dendritic cells^68^. We identified a novel mechanism of immunosuppression mediated by the T3SS that target eosinophils, and possibly other immune cells, such as epithelial cells, to promote secretion of IL1RA. Overall, our findings demonstrate that bacteria utilize the T3SS to promote expression of IL1RA which then enhance colonization and long-term persistence. Treatment with anti-IL1RA antibodies decrease bacterial burden in the lungs, providing novel avenues for therapeutic development. These results provide some insights into the side effects of long-term treatment with anakinra. Moreover, our results also highlight the need of a tight balance of the IL1 signaling axis which is required to prevent the auto-immune disorders, while allowing for the clearance of inflammatory diseases. Finally, IL1RA is also increased in fungal triggered asthma^29^, which might further support that recurrent infections^69^ could promote a chronic increase in IL1RA. This constant increase in IL1RA might reach a point from where it cannot be controlled anymore, leading to the development of asthma in infants.

## MATERIALS AND METHODS

### Bacterial Strains and Culture Conditions

*B. bronchiseptica* strain RB50, a *btrS* knockout mutant^10^, and a *bscN* knockout mutant^38^ were used in this study. *B. bronchiseptica* RB50 and mutants were cultured Difco Bordet-Gengou (BG) agar (BD, cat. 248200) and supplemented with 10% sheep defibrinated blood with 20 μg/mL streptomycin or classical LB broth as previously described^7^. Our *B. pertussis* strain BP536 and *B. parapertussis* BPP12822, were grow in BG agar or Stainer-Scholte as previously described^7^.

### Animal Experiments

Our animal experiments included Balb/c and il1rn^-/-^ mice, mice originally purchased from Jackson laboratories, Bar Harbor, ME, and then bred in our facilities. Our breeding colonies were kept under the care of the employees and veterinarians of Louisiana State University Health - Shreveport Animal Care Facility, Shreveport, LA, (AUP:20-038, P22-031). All experiments were carried out in accordance with all institutional guidelines (AUP:20-0038, P22-031).

For our animal inoculations mice were anesthetized with 5% isoflurane and when sleep they were intranasally challenged with 30-50μl of PBS containing 1×10^5^ CFU/mL *B. bronchiseptica* RB50, RB50Δ*bscN* and RB50Δ*btrS*.

Anti-IL1RA treatment was performed daily. Mice were anesthetized with 5% isoflurane and when asleep, they are intranasally inoculated with 10μl of 500 μg of anti-IL1RA antibodies purchased from Leinco technologies (Anti Mouse/Human IL-1ra/IL-1F3 – 500 μg). We started the treatment at three different time points, day 1 post-infection, day 3 post-infection and day 5 post-infection. Supplementation with IL1RA was performed daily from day 1 post-infection by intraperitoneally delivering 25 μg of murine IL1RA purchased from sigma (SRP6006-50UG). Mice were euthanized using 5% CO_2_ followed by cervical dislocation. For all animal protocols we followed the guidelines of AALAC, and the protocols described on our approved IACUC P20-038 and P-22-031.

### Mouse transcriptomics

At day 7 post-inoculation, mice were euthanized according with humane endpoints and our protocol approval. Lungs were collected in 4 different beads tubes containing Trizol and kept in ice for the shortest time possible before being homogenized and freeze at -20°C. For RNA extraction, we follow the protocols recommended by the manufacturer, PureLink RNA Extraction kit, Invitrogen, and the 4 different tubes were pull together to have the whole lung RNA contained in one only preparation.

Total RNA was extracted using the PureLink RNA Mini Kit (Invitrogen, Waltham, MA, United States) and treated with PureLink DNase (Invitrogen, Waltham, MA, United States) following manufacture protocols. RNA concentrations were quantified using a Nanodrop One (ThermoFisher Scientific, Waltham, MA, United States)^7^.

One μg of RNA was used for qRT PCR. To perform the qRT PCR we followed the recommendations of the manufacturer Luna Universal One-Step qRT-PCR Kit (New England BioLabs Inc., Ipswich MA, United States qRT PCR reactions were carried out in the Genomics Core Facility at Louisiana State University Health – Shreveport on a BIO-RAD CFX96 (LSU Health – Shreveport, LA, United States). Primer sequences are shown in table 1.

### Immunohistochemistry

For all pathology experiments, mice were euthanized and perfused intratracheally with sterile PBS and 4% paraformaldehyde (PFA). Lungs were subsequently fixed overnight in 10 mL of 4% PFA prior to processing and paraffin embedding. Lungs are processed, paraffin-embedded, sectioned in .5 μm thick slices, and placed on glass slides. Slides with PFA fixed and paraffin-embedded tissues were deparaffinized in consecutive washes in xylene, followed by rehydration in decreasing concentrations of ethanol ranging from 100% - 50%. Staining was performed following previously published methods^70^. Immunofluorescent images of the lungs were captured using a Keyence BZX-800 microscope. Image analysis was conducted using the Keyence Image Analysis Software.

### Statistical analysis

All experiments were performed in three independent biological replicates. The exact number of mice and technical replicates is indicated in each figure legend. For animal experiments we performed Two-Way ANOVA analysis, while qRT-PCR experiments were analyzed using One-Way ANOVA analysis. All graphs and data were analyzed using the GraphPad V9.0. A p value Δ 0.05 was considered statistically significant. In the figures the asterisks correspond with * ≤0.05, ** ≤0.01, *** ≤0.001, and **** ≤0.0001.

## ACKNOWLEDGEMET

We would like to acknowledge the support of the funding bodies: NIH COBRE award 1P20-GM134974-0181. Louisiana Board of Regents LEQSF(2022-25)-RD-A-33. Center of Excellence for Arthritis and Rheumatology intramural award. Intramural Research Council Seed package support from LSU Health, Shreveport. Start-up package from LSU Health, Shreveport.

We would also like to acknowledge Martin Sapp for their ideas, brainstorming, and support during the editing process. We would like to thank Dr. Matthew Woolard for the brainstorming and ideas. We would like to acknowledge the SMART program at Louisiana State University as one of the authors, Sarah Johnson is part of this program.

## Author contribution

TW performed in vitro experiments, analyzed data, contributed to writing and editing. CR performed in vivo and in vitro experiments, analyzed data, contributed to the editing. NF performed in vivo experiments, analyzed data, contributed to writing and editing. SJ performed in vitro experiments, analyzed data, and contributed to writing and editing. MCG conceptualized the project, designed experiments, performed experiments, analyzed data, writing and editing, obtained funding, supervised.

